# Molecular chaperones maximize the native state yield per unit time by driving substrates out of equilibrium

**DOI:** 10.1101/153478

**Authors:** Shaon Chakrabarti, Changbong Hyeon, Xiang Ye, George H. Lorimer, D. Thirumalai

## Abstract

Molecular chaperones have evolved to facilitate folding of proteins and RNA *in vivo* where spontaneous self-assembly is sometimes prohibited. Folding of *Tetrahymena* ribozyme, assisted by the RNA chaperone CYT-19, surprisingly shows that at physiological Mg^2+^ ion concentrations, increasing the chaperone concentration reduces the yield of native ribozymes. In contrast, the more extensively investigated protein chaperone GroEL works in exactly the opposite manner—the yield of native substrate increases with the increase in chaperone concentration. Thus, the puzzling observation on the assisted-ribozyme folding seems to contradict the expectation that a molecular chaperone acts as an efficient annealing machine. We suggest a resolution to this apparently paradoxical behavior by developing a minimal stochastic model that captures the essence of the Iterative Annealing Mechanism (IAM), providing a unified description of chaperone mediated-folding of proteins and RNA. Our theory provides a general relation involving the kinetic rates of the system, which quantitatively predicts how the yield of native state depends on chaperone concentration. By carefully analyzing a host of experimental data on *Tetrahymena* (and its mutants) as well as the protein Rubisco and Malate Dehydrogenase, we show that although the absolute yield of native states decreases in the ribozyme, the rate of native state production increases in both the cases. By utilizing energy from ATP hydrolysis, both CYT-19 and GroEL drive their substrate concentrations far out of equilibrium, in an endeavor to maximize the native yield in a short time. Our findings are consistent with the general expectation that proteins or RNA need to be folded by the cellular machinery on biologically relevant timescales, even if the final yield is lower than what equilibrium thermodynamics would dictate. Besides establishing the IAM as the basis for functions of RNA and protein chaperones, our work shows that cellular copy numbers have been adjusted to optimize the rate of native state production of the folded states of RNA and proteins under physiological conditions.

**Significance statement:** Molecular chaperones have evolved to assist the folding of proteins and RNA, thus avoiding the deleterious consequences of misfolding. Thus, it is expected that increasing chaperone concentration should lead to an enhancement in native yield. While this has been observed in GroEL-mediated protein folding, experiments on *Tetrahymena* ribozyme folding assisted by CYT-19, surprisingly show the opposite trend. Here, we reconcile these divergent experimental observations by developing a unified stochastic model of chaperone assisted protein and RNA folding. We show that chaperones drive their substrates out of equilibrium, and in the process maximize the rate of native substrate production rather than the absolute yield or the folding rate. *In vivo* the number of chaperones is regulated to optimize their functions.

## I. INTRODUCTION

Small single domain proteins fold rapidly with sufficient yield as envisioned by Anfinsen[1–3]. However, larger proteins, especially those with complex native state topology, are often kinetically trapped in metastable states for sufficiently long times that protein aggregation could be a major problem [4–6]. In such non-permissive conditions, the native state yield is extremely low. To remedy this deleterious situation, molecular chaperones have evolved to rescue those substrate proteins (SPs) that are prone to aggregate, and hence do not fold spontaneously with enough yield on cellular time scales [7, 8]. The best studied example is the bacterial chaperonin GroEL [8, 9] a promiscuous stochastic ATP-consuming machine that mediates folding of a variety of SPs regardless of the topology they adopt in the native state. Indeed, nearly thirty years ago it was shown that GroEL recognizes SPs as long as they are in the misfolded state, thus exposing hydrophobic residues which are usually buried in the native state [10]. In contrast to protein chaperones, much less is known about RNA chaperones and their functions. Under typical *in vitro* conditions, ribozymes readily misfold into a manifold of metastable states [11–13]. The barriers between these states and the native state is much larger than the thermal energy, thus making the time scales for transition between the metastable states and the native state longer than the cell doubling time.

The need for chaperones is best understood by considering spontaneous folding of proteins and RNA folding under non-permissible conditions. The kinetic partitioning mechanism (KPM) provides a common unifying description of chaperone-free folding of SPs as well as ribozymes in rugged energy landscapes [11, 14–16]. According to the KPM (illustrated in Fig. 1), a fraction of the initial population of molecules, referred to as the partition factor Φ, folds rapidly to the native state while the remaining fraction, (1 – Φ), is kinetically trapped in misfolded states for times exceeding viable biological times. Proteins and ribozymes, which reach the folded state with sufficient yield (Φ ~ 1) do not require chaperones. In contrast, the non-bacterial protein Rubisco with(Φ < 5% [17] and *Tetrahymena* ribozyme (Φ ~ 8 %) [11–13] require assistance from chaperones.

**FIG. 1:**
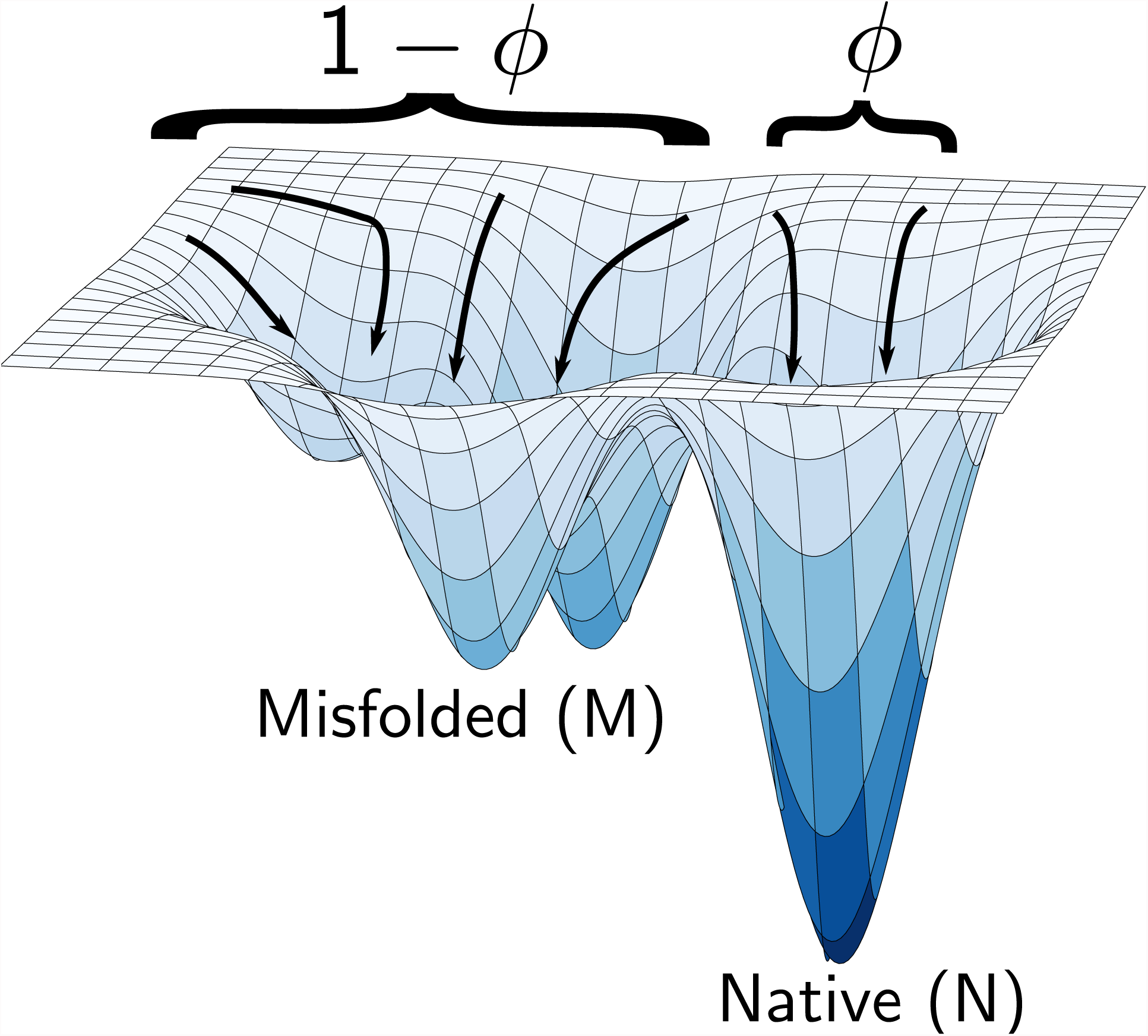
Illustrating the kinetic partitioning mechanism of spontaneous folding of proteins and RNA in a rugged folding free energy landscape. Under non-permissive conditions, only a fraction Φ of the unfolded molecules fold rapidly to the native state. The rest (1 – Φ)) misfold and remain kinetically trapped in a manifold of long-lived metastable states. If Φ is small, as is the case in the folding of ribozymes and proteins under non-permissible conditions, their folding requires the chaperone machinery.

The mechanism of GroEL action has mysteriously remained controversial. Two diametrically opposite models have been proposed — the passive or active Anfinsen cage mechanism, which posits that the GroEL encapsulates the misfolded SP and provides an environment that avoids SP-SP interaction, thereby preventing aggregation and promoting folding [18, 19]. It should be noted folding rate could be enhanced due to confinement in a non-cycling artificial GroEL mutant [20], as predicted theoretically [21–23]. On the other hand, the Iterative Annealing Mechanism (IAM) proposes that GroEL utilizes the energy from ATP binding and hydrolysis, leading to large conformational changes in GroEL [17, 24–26]. The allosteric changes lead to partial unfolding and encapsulation of the misfolded SP, thus giving the SP another chance to fold to the correct native state. Iterating the cycles of encapsulation and unfolding eventually leads to a high yield of the native SPs [17]. Unlike the cage model the IAM quantitatively explains all of the available experimental data, including the effect of GroEL mutants in the rescue of mitochodrial Malate Dehydrogenase (mtMDH) and Citrate Synthase [27].

In the context of chaperone mediated RNA folding, a large number of studies unequivocally suggest that the IAM is the dominant mechanism of RNA folding [28–30] without the benefit of quantitative analyses. This has been attributed to the ubiquitous role of superfamily-2 (SF2) helicases performing RNA re-modeling and chaperone activities, many of which are known to unwind nucleic acids either processively or in a local manner [31, 32]. In RNA, therefore, it seems to be clear that kinetic traps are resolved by first partially unwinding (thereby unfolding) misfolded states of the substrate.

The proposal that assisted folding by GroEL and RNA chaperones involves unfolding of the substrates suggests that there ought to be a universal mechanism of chaperon-assisted protein and RNA folding, which we referred to as the generalized iterative annealing mechanism in an earlier work [33]. The generalization is required since it was shown, that unlike the case of proteins where chaperones recognize only the misfolded SPs, the RNA chaperones also bind and unwind native RNA folds [28, 29]. Therefore, while the basic principle of repeated rounds of misfolded substrate recognition and partial unfolding remains identical between proteins and RNA chaperones, there is an additional element of native state recognition by the chaperone in case of RNA. However, this additional aspect of native state recognition leads to an immediate conundrum, which was observed in experiments of CYT-19 mediated folding of *T.* ribozymes — as the chaperone (in this case CYT-19) concentration is increased, the final yield of native state is reduced [28]. This finding is qualitatively different from the observations on GroEL-mediated folding of proteins, where increasing the chaperone concentration increased the final yield of the native protein [17].

Here, we create a stochastic model based on the generalized IAM, which quantitatively explains the apparently paradoxical results in the function of RNA and GroEL within a unified theory. Using our theory, we analyze a host of experimental data on CYT19-mediated folding of the *Tetrahymena* ribozyme (and its mutants) as well as Rubisco, and recent data on mtMDH [34]. In both cases, the *rate* of increase of the native state yield increases with increasing chaperone concentration. Therefore, our work shows that both the RNA and protein chaperones have evolved to maximize production of the native state per unit time, and not simply the absolute yield or the folding rate. We further show that the chaperones achieve optimal performance by hydrolyzing ATP and driving their substrates into steady states that are far from equilibrium. While equilibrium thermodynamics would predict far higher long term native yields in assisted folding of proteins and RNA, cellular time scales are much shorter than the times needed to reach equilibrium. Our work, therefore, suggests that cells settle for less native substrate than is theoretically possible, but obtained over much faster time scales by allowing chaperones to utilize non-equilibrium processes.

## II. UNIFIED MODEL FOR PROTEIN AND RNA FOLDING BY CHAPERONES

The crucial prediction of the IAM for proteins [17] is that GroEL binds to misfolded substrate proteins and unfolds it fully or partially, giving the SP another chance to fold correctly. RNA chaperones (CYT19 for example) are more indiscriminate in that they also unfold the native substrate [28]. We subsume these scenarios using a three-state model, which we demonstrate not only fits all the data but leads to general principles of chaperone function, and the associated optimization problem that nature has solved to prevent aggregation. We define the three main states I, N and M, corresponding to Intermediate, Native and Misfolded respectively (Fig. 2). The substrate may be fully or partially unfolded by the chaperone [8, 28]. Hence, the free substrates, prior to folding, belong to the I state in our model. We assume that the chaperones do not bind to the unstructured I state because doing so would result in an unstable complex. In addition, to describe the action the chaperone concentration dependence on folding rates, we define two additional states CM and CN, corresponding to chaperone bound to misfolded and native states, respectively (Fig. 2).

**FIG. 2:**
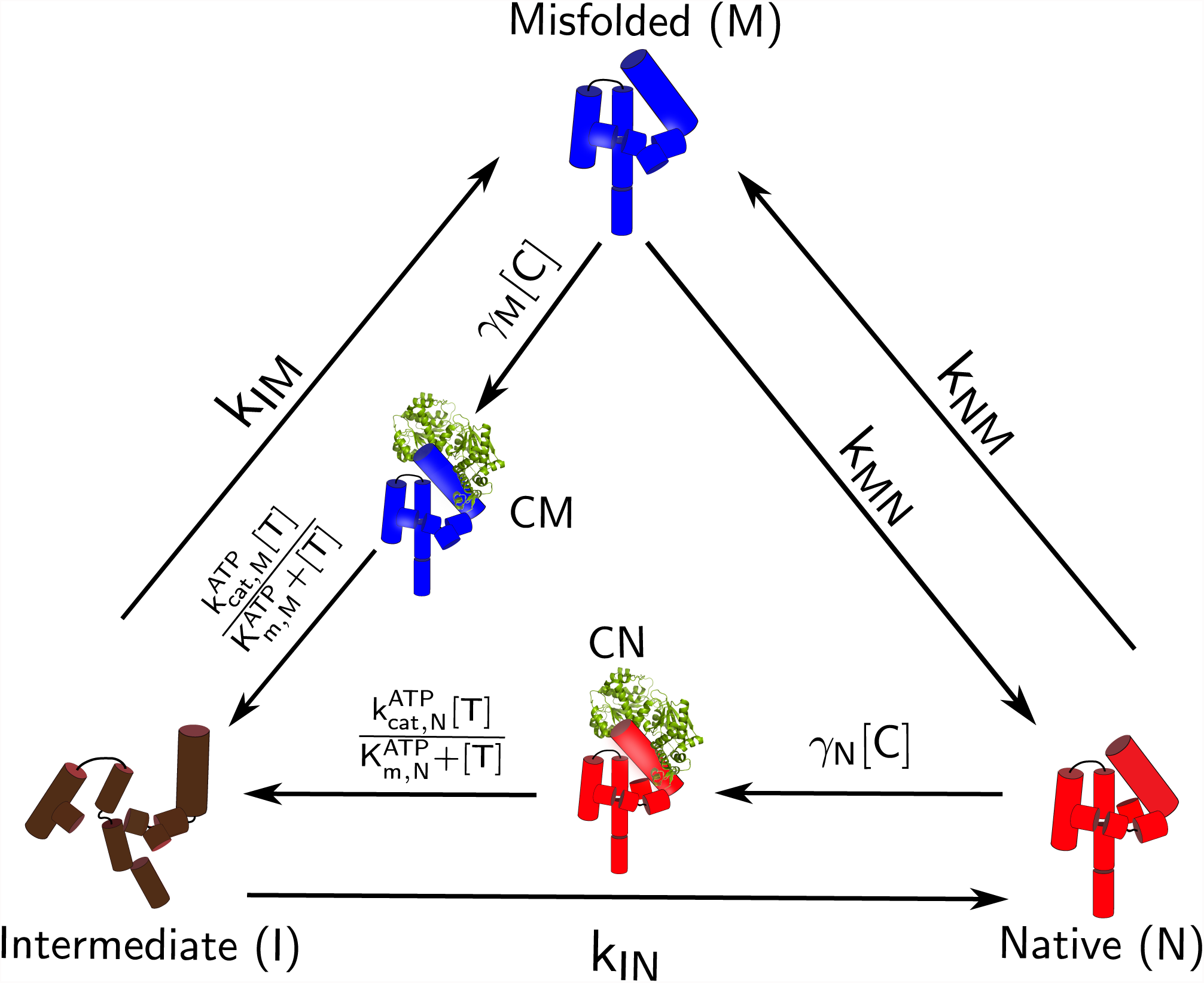
Schematic illustration of the model for a unified description of chaperone assisted folding. We use *Tetrahymena* ribozyme for illustration purposes. The ribozyme in the intermediate (I, brown), native (N, red), misfolded (M, blue) states and RNP complexes are illustrated in the scheme. The CYT-19 is represented in green. The same model is applicable to describe GroEL-associated folding except CYT-19 is replaced by the chaperonin machinery.

The rates *k_ij_* correspond to transitions from state *i* to *j,* where *i, j* = *I, N, M.* Starting from an ensemble of SPs in the unfolded state, the ribozyme population (or SPs) rapidly collapses to the intermediate [11]. From the I state, a fraction of molecules (Φ) fold rapidly to the native state N while the rest of them (1 – Φ) are trapped in long-lived metastable misfolded intermediates M, which slowly fold to the native state, as predicted by the KPM [11, 35, 36]. The kinetic partition factor Φ is related to *k*_IN_ and *k*_IM_ by

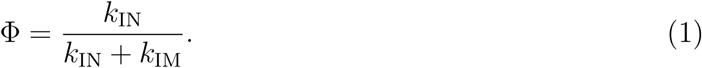

Although there are a multitude of states in the intermediate and misfolded ensembles in a rugged folding landscape [36], we subsume all such possible states into I and M for simplicity. *In vitro* experiments, in the absence of chaperone, suggest a rapid equilibration between the multitude of states in the misfolded ensemble, and hence the misfolded to native state transition can be described by a single rate [11, 35, 37, 38]. The effective rates *k*_IN_ and *k*_IM_ are assigned to the transitions to the native and misfolded states, respectively.

To model the function of the GroEL and Cyt-19, we allow the chaperone to recognize both M and N, taking the misfolded and native states back to the intermediate state I with rates *k*_MI_ and *k*_NI_, respectively. The chaperone–ribozyme or chaperone–protein complexes formed with misfolded and native states, denoted as CM and CN respectively, utilize energy obtained from ATP hydrolysis to revert to the partially unfolded intermediate I.

The chaperone binds to M and N with second order rate constants *γ_M_* and *γ_N_,* respectively. Note that *γ_N_* and *γ_M_* can be interpreted as the effective rate of protein/ribozyme recognition and processing by the chaperone, which reduces to *k*_cat_/*K*_M_ at low chaperone concentration, assuming that GroEL/CYT-19 binding to M or N follows Michaelis-Menten kinetics [29]. It has been shown that the CYT-19 dependent rate of unfolding of ribozyme is linear up to 500 nM, indicating that the lower bound of the effective Michaelis-Menten constant 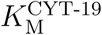 is 500 nM. We assume that the transitions from CN or CM to I follow Michaelis-Menten kinetics with the ATP concentration [*T*], with distinct turnover rates 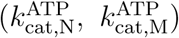, and Michaelis constants 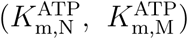 (Fig. 2). The transition rates involving the states CM and CN, shown in Fig.2, can be subsumed into one by observing that each of the transition times for M→ I and N→ I is a sum of two time scales, i.e., binding of chaperone to substrate and ATP-dependent partial unfolding. Therefore, the overall rates of chaperone and ATP mediated unfolding rates can approximately be written as 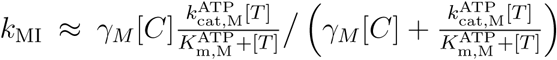 and 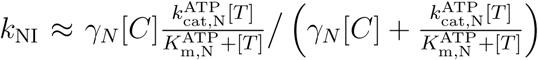. We are thus left with only the three states I, M and N, which greatly simplify the analyses of the experimental data. Finally, the transition rate from M to N is *k*_MN_ while the reverse rate is kNM, such that the free energy difference between the two states is given by Δ*G*_NM_ = *G*_M_ – *G*_N_ = –*k_B_T*log(*k*_NM_/*k*_MN_).

## III. RESULTS AND DISCUSSION

### Assisted kinetics and the native state yield of proteins and RNA

The time evolution of the probability of the system being in the state *i* at time *t P*(*i, t*) is given by the master equation,

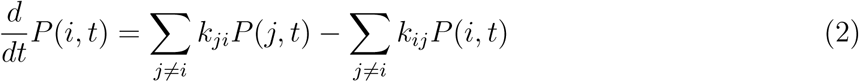

with *i, j* = I, N, M. The non–equilibrium steady-state yield population of the native state *P*(*N, t* → ∞), is reached from any initial condition, and is given by, (see **SI** for details):

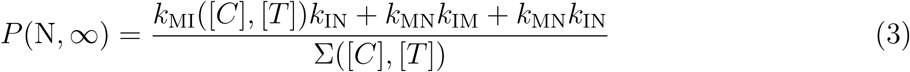

where ∑([*C*], [*T*]) = *k*_NI_([*C*], [*T*])*k*_MN_ + *k*_IM_*k*_NI_([*C*], [*T*]) + *k*_IM_*k*_MN_ + *k*_IN_*k*_NM_ + *k*_NM_*k*_MI_([*C*], [*T*]) + *k*_MI_([*C*], [*T*])*k*_IN_ + *k*_IM_*k*_NM_ + *k*_MI_*k*_NI_([*C*], [*T*]) + *k*_MN_*k*_IN_. Note that this steady-state value of the native yield depends on both the concentration of the chaperone [C] as well as the ATP concentration [T]. If *k*_NI_ = *k*_MI_ = 0, which happens when either [C]=0 and/or [T]=0, the native population becomes,

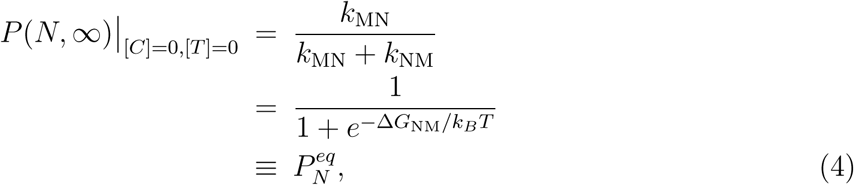

which is the expected result at thermodynamic equilibrium (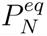) between the two states, N and M.

Remarkably, the behavior of the steady state native substrate yield *P*(*N*, ∞) under the action of RNA chaperone CYT-19 is strikingly different compared to that of GroEL. While the steady state yield of the folded protein *increases* on increasing the GroEL concentration, the steady state yield of native ribozyme *decreases* on increasing CYT-19 concentration. The contrasting behavior is fully explained by our model. Our theory predicts that *P*(N, ∞) monotonically increasing function of [C], if the inequality,

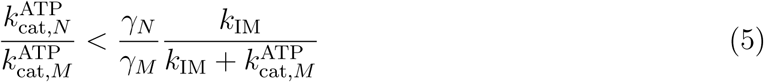

holds. On the other hand, if

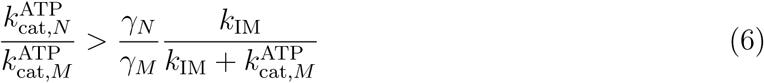

then *P*(N, ∞) will be a monotonically decreasing function of [C] (see **SI** for details).

Substituting the parameters from Table S1 (see sections below), we see that the inequality in Eq. 6 is indeed satisfied by *T.* ribozyme. Similarly, using the best-fit parameters from Table S2 shows that Rubisco satisfies the inequality in Eq. 5, thus explaining the increase in native yield of Rubisco as GroEL concentration is increased.

### Generalized IAM and the native state recognition factor

Without chaperones, only a small fraction Φ of the original unfolded ensemble reach the native state spontaneously. The rest, 1 – Φ remain trapped in long-lived metastable states. In order to rescue these kinetically trapped proteins to the native state, the chaperone molecules recognize and bind to the exposed hydrophobic regions of the misfolded protein. The entire fraction 1 – Φ is then assisted by GroEL, in all likelihood reverting it to the more expanded form, and the whole process is repeated over and over again. The yield of native state as a function of such reaction cycles *n,* is given by *Y_N_*(*n*) = 1 – (1 – Φ)*^n^*. As *n* becomes large, the native yield can theoretically reach *Y_N_*(*n*) → 1.

The generalized IAM [33] allows for the possibility of native state recognition by the RNA chaperone, CYT-19, which was not considered previously [39]. The chaperone is allowed to act on the native state in addition to the misfolded states of protein or RNA, and redistributes *κ*Φ again into *κ*Φ^2^ native states and *κ*Φ(1 – Φ) misfolded states, where *κ* (0 < *κ* < 1) is the degree of discrimination of by the chaperone between the native and misfolded state. A fraction (1 – *κ*)Φ of the original native population remains unperturbed in the same native state. It is easy to show that, the net gain in the fraction of native state after *n* iterations is given by Φ((1 – Φ)(1 – *κ*))*^n^*^−1^ (where *n* = 1, 2,…). The total yield of the native state after *n* iterations *Y_N_*(*n*), is therefore *Y_N_*(*n*) = Φ + Φ(1 – Φ)(1 – *κ*)+…+(Φ((1 – Φ)(1 – *κ*))*^n^*^−1^, and the conditions of Φ < 1 and *κ* < 1 lead to,

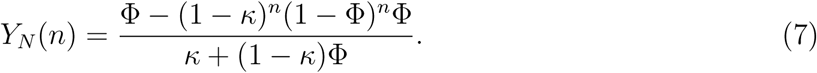

The physical meaning of the discrimination factor, *κ*, is evident, by making an approximate mapping of the long-time yield *Y_N_*(*n* → ∞) to the equivalent expression in our master–equation framework, *P*(N, ∞). By substituting 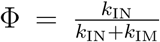 into Eq.7 and taking the limit *n* → ∞ we obtain 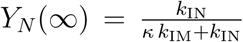, while *P*(N, ∞) with *k_MN_*, *k_NM_* ≪ 1 and *k_NI_* ≪ *k_IN_* reduces to 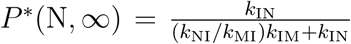. Therefore, *κ* is approximately the ratio of two rate constants associated with chaperonin induced unfolding:

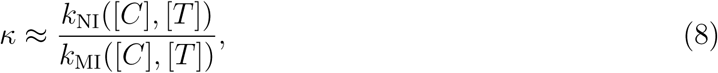

which is in accord with our intuitive definition of *κ* given in Eq.7. It is worth noting that *κ* depends on the chaperone ([*C*]) and ATP ([*T*]) concentrations, which suggests that it is possible to reduce *κ* by increasing [*C*] or [*T*]. Evidently, for GroEL *κ* → 0 because *k*_NI_ ([*C*], [*T*]) is negligible.

### CYT-19 mediated folding of *T.* ribozymes

Since the discovery of self-splicing enzymatic activity in the group I intron *Tetrahymena thermophila* ribozyme [40–42], the *T.* ribozyme has been the workhorse used to reveal the general principles of RNA folding. In accord with the KPM (Fig. 1), the value of Φ of the wild-type *T.* ribozyme that attains catalytic activity in the absence of CYT-19 is only (6–10) % (at 25 °C), while the majority of ribozymes remain inactive [11, 13]. In the case of *T.* ribozyme, it is suspected that the premature formation of the incorrect base pairs stabilize the misfolded conformations[43]. For example, to disrupt a six base-paired helix, a secondary structure motif ubiquitous in RNA, the free energy barrier is *δG*^‡^ =10–15 kcal/mol (= 5 stacks x (2–3) kcal/mol/stack). The timescale, 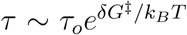 with *τ_o_* ≈ 1 *μ*sec [44], for a spontaneous disruption of base stacks is estimated to be 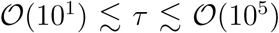 sec. Thus, once trapped into a mispaired conformation it is highly unlikely to autonomously resolve the kinetic trap on a biologically viable timescale [43].

We first analyze the rate of CYT-19 in facilitating the folding of *T.* ribozyme. Time resolved kinetics of two variants (P5a mutant and P5abc-deleted (ΔP5abc) ribozyme) as well as the wild-type (WT) of the ribozyme were probed by varying CYT-19 and ATP concentrations [28]. We establish the validity of our theory by using Eq. S2 in the SI to quantitatively fit an array of experimental data on the WT and P5a mutant (Figs 3a and 4a-b respectively). In the experiments, the fraction of native ribozyme was probed as a function of time, under different initial conditions: (i) starting from completely folded (N) ribozymes; (ii) starting from primarily misfolded (M) ribozymes; (iii) CYT-19 chaperone inactivated by addition of proteinase K. To probe the effects of CYT-19 and ATP on the production of active (native) state, CYT-19 was varied for the cases (i) and (ii), and ATP concentration was varied for the case (i). In total, we used our theory to fit five sets of data for the WT (Fig. 3a) and eleven sets of data for the P5a variant (Fig. 4a-b) ribozyme. By accounting quantitatively for the data set we extracted the best fit parameters, given in Table S1.

**FIG. 3:**
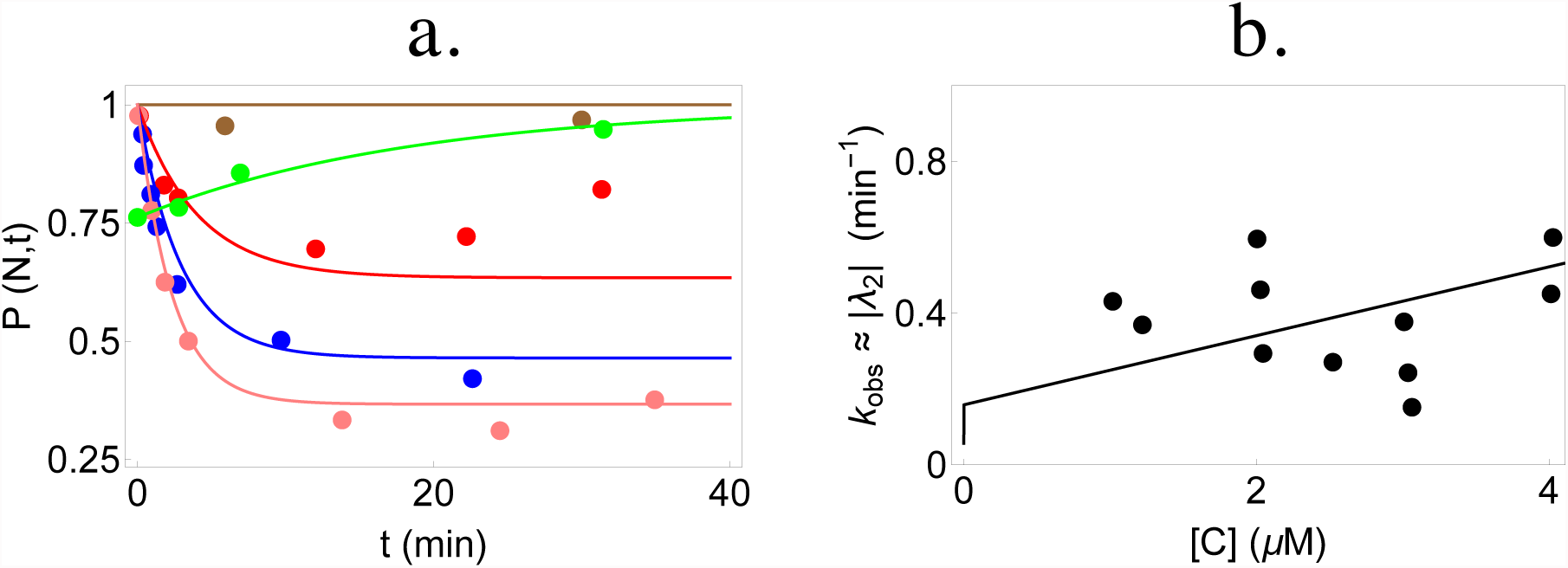
Analysis of data on the wild-type (WT) *Tetrahymena* ribozyme. Circles represent experimental data while the curves are plots of Eq. S2. The five sets of data in **a** have been fit simultaneously to determine the best parameters for the WT (given in Table S1). (**a**). Kinetics of WT ribozyme in 2 mM ATP concentration for various concentrations of CYT-19: no CYT-19 (brown), 1 *μ*M CYT-19 (red), 2 *μ*M CYT-19 (blue) and 3 *μ*M CYT-19 (pink). The curve in green is obtained for a mixture of native and misfolded WT ribozyme when proteinase K is introduced to inactivate CYT-19. (**b**). Dependence of *k_obs_* of WT ribozyme on CYT-19 (data from Fig. 1d of [28]). The curve is the CYT-19 dependence of the second eigenvalue |λ_2_| obtained from our model (see SI), with parameters obtained from the fits in **a.** Given the large experimental uncertainty only the trend, *k_obs_* increases as [C] increases, is meaningful.

The overall trends in the parameters, extracted by simultaneous fit of the available data, are consistent with the direct experimental measurements and estimates (see Table S1). Note that some of the experimental results cited in Table S1 were performed under different conditions (temperature, Mg^2+^ ion concentration or absence of CYT-19) than the experiments analyzed using our theory. These differences could affect the various rates and are pointed out in Table S1. For the P5a mutant, the fraction Φ of ribozymes that fold directly to the native state, was estimated to be 0.09 [28] while Φ (Eq.1) calculated from our fitted values of *k*_IM_ and *k*_IN_ is 0.10 – 0.12. The free energy difference, Δ*G*_NM_, calculated from *k*_NM_ and *k*_MN_ gives 2.6 *k_B_T.* This value is in rough accord with experimental results showing that the native state of P5a is less stable compared to the WT with 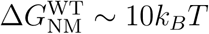 [11, 45].

For all three variants of the ribozyme, one dominant eigenvalue (|λ_2_|) of the master equation formulation describes the overall kinetic behavior of the three state model (see SI). Thus, the time evolution of the fraction of native state is primarily governed by the exponential term 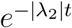, making **|**λ_2_**|** comparable to the experimentally observed rate, *k_obs._* In order to assess the effect of varying ATP and CYT-19 concentration on the chaperone-induced unfolding kinetics of the native ribozyme, we compared |λ_2_**|** (computed from the parameters in Table S1) with data on *k_obs_* as a function of CYT-19 (Fig. 3b for WT, Fig. 4c for P5a) and ATP concentration (Fig. 4d for P5a). The reasonable agreement of these curves with the experimental data and the best-fit parameters with their corresponding experimentally measured values, indicates that our kinetic model faithfully describes CYT-19 mediated folding/unfolding of *T.* ribozyme. The agreement is especially satisfactory given the large scatter in the experimental data.

**FIG. 4:**
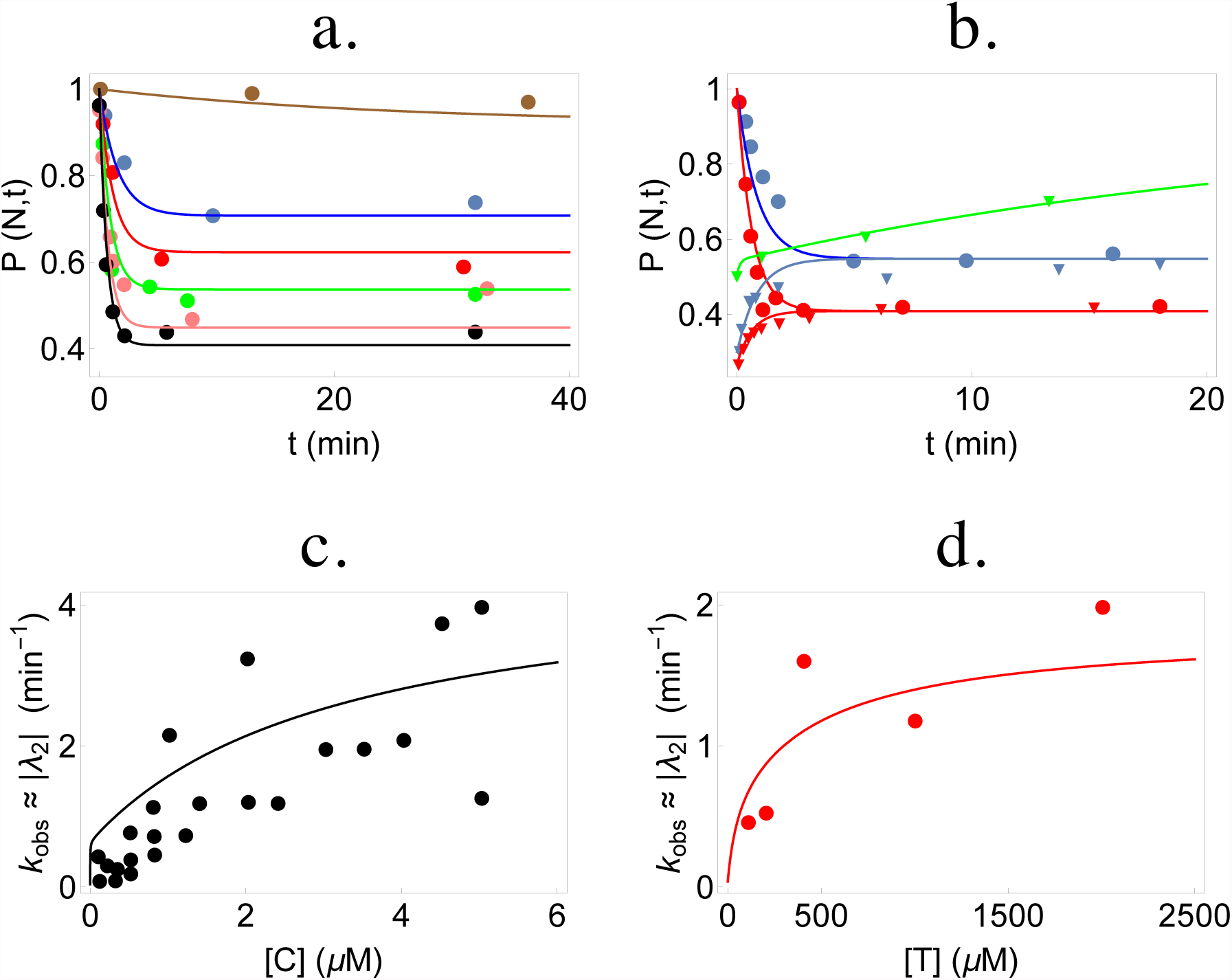
Analysis of experimental data on the P5a variant of *Tetrahymena* ribozyme. The circles and inverted triangles represent experimental data while the curves are plots of Eq. S2. The eleven sets of data in **a-b** were fit simultaneously to determine best-fit parameters (given in Table S1). (**a.**) CYT-19 (1 *μ*M)-induced kinetics starting from the native P5a variant ribozyme in 5 mM-Mg^2+^ at various ATP concentrations: no ATP (brown), 100 *μ*M-ATP (blue), 200 *μ*M-ATP (red), 400 *μ*M-ATP (green), 1 mM-ATP (pink), 2mM ATP (black). (**b**.)(**c-d**). Dependence of *k_obs_* of P5a variant on CYT-19 (data from Fig. S3 of [28]) and ATP concentration (data from Fig. S4c of [28]) respectively. The lines are CYT-19 or ATP dependence of the second eigenvalue |λ_2_| obtained from our model, with parameters obtained from the fits in **a-b.**

The ratio, *k*_MI_/*k*_NI,_ which quantifies how indiscriminately the chaperone unwinds both the native and misfolded states, is roughly 40-80, in the ribozyme 0.5 – 5*μ*M concentration range of CYT-19 and at 1 mM ATP using the parameters for the P5a variant in Table S1. We obtained qualitatively similar results if parameters from the WT are used. Since more of the P5a parameters could be robustly fit, we report *k*_MI_/*k*_NI_ for only the P5a variant.

Finally, to test the importance of the native state recognition by CYT-19, we analyzed how the long term native state yield (Eq. 3) changes due to perturbations of the parameter γ*_N_* around the best fit values of the WT ribozyme (Fig. S2). We also perturbed some of the other parameters that could conceivably be changed by making mutations in the chaperone domains, for example the ATP hydrolysis rates and the binding constant γ*_M_* (Fig. S2). Interestingly, *P*(*N*, ∞) is most sensitive to changes in γ*_N_* as compared to the other parameters (Fig. S2), thereby indicating that changes in recognition and binding of CYT-19 to native RNA can result in significant shifts in the final native state yield.

### GroEL mediated folding of Rubisco and MDH

Rubisco is a stringent substrate for GroEL in the sense that the full machinery including ATP and GroES is required to ensure folding. In a previous study, the GroEL-assisted folding of Rubisco as a function of GroEL concentration was reported [17]. Starting from acid denatured Rubisco (in kinetically trapped misfolded states), the yield of the native state increased with time upon addition of the chaperonin system (GroEL and GroES). Using our theory, we simultaneously fit the nine time-evolution curves, corresponding to nine different concentrations of GroEL using Eq. S2. From the results in Fig. 5, we draw the following general conclusions: (i) The excellent fit in Fig. 5a shows that the model quantitatively captures the kinetics of GroEL-assisted folding of Rubisco; (ii) The folding rate increases non-linearly with GroEL concentration; (iii) Most importantly, the dependence of *P*(*N, t* ≈ 60mins) on chaperonin concentration shows that equilibrium is not reached at long times, despite GroEL being in far excess of the concentration of Rubisco. This crucial point, which also holds for CYT-19 assisted folding of *T.* ribozyme, is further discussed below.

**FIG. 5:**
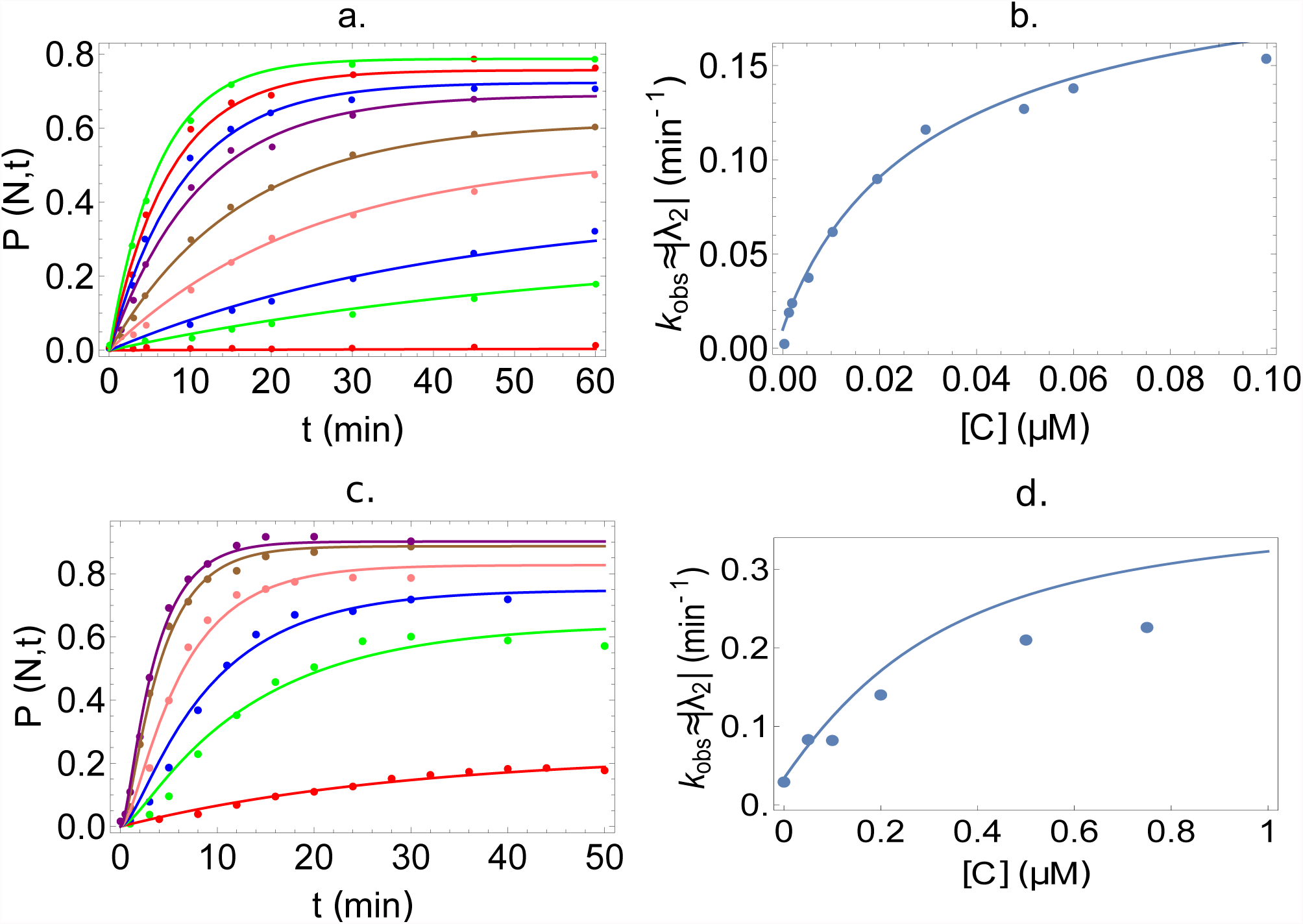
Using the kinetic model to analyze GroEL-assisted folding of Rubisco and MDH. (**a**) Quantitative fits of time-dependent increase of folded Rubisco at various GroEL concentrations and 1mM ATP, using our kinetic model (data taken from [17]). Note that the y-axis was converted to a fraction between 0 and 1 by dividing Rubisco concentrations by 50 nM, which was the starting concentration of acid-denatured protein in [17]. The concentrations of GroEL are (from bottom to top): 0, 1, 2, 5, 10, 20, 30, 50 and 100 nM. A single set of parameters, listed in Table S2, fits all the curves. The dependence in the steady state values of *P*(*N, t*) (*t* ≈ 60 min) is an indication of the departure from equilibrium. In all cases, *P*(*N, t*) (*t* ≈ 60 min is less than the equilibrium value. (**b**) Dependence of the rate *k_obs_* as a function of GroEL concentration for folding of Rubisco. (**c**) Fits of time-dependent increase of native MDH at various GroEL concentrations and 500*μ*M ATP. Concentrations of GroEL are (from bottom to top): 0, 0.05, 0.1, 0.2, 0.5 and 0.75 *μM.* (**d**) Dependence of the rate *k_obs_* as a function of GroEL concentration for folding of MDH. In all panels the dots represent data from experiments while the curves represent theoretical predictions.

The fitted parameters for GroEL mediated folding of Rubisco (shown in Table S2) are consistent with previous experimental and theoretical results. From our fits, we obtain the partition factor Φ ~ 0.01, remarkably close to the expected value from experiments and a previous theory [17]. The binding rate of GroEL was measured to be 10^7^ – 10^8^ *M*^−1^*s*^−1^ [46] while we obtain γ*_M_* ≈ 0.7 x 10^7^*M*^−1^*s*^−1^. A previous kinetic model was developed by Tehver and Thirumalai, to describe the coupling between allosteric transitions of GroEL during the folding of Rubisco [27]. Though the details of the model are completely different from the present work, we can compare some results of both the models. Our estimate of the GroEL binding rate is consistent with the value of 10^7^*M*^−1^*s*^−1^, and Φ, extracted from their fits was 0.02. The consistently small value of the partition factor Φ( ≈ 0.01 – 0.02) implies that in a cycling system, which is most relevant for *in vivo* function of GroEL, only about (1–2) % of Rubisco reaches the folded state in the absence of chaperone.

We should point out that unlike the ribozyme analysis, there is not enough data in the GroEL-Rubisco experiment to extract *k*_NM_ and *k*_MN_ (and henceΔ*G*_NM_) accurately. However, this does not affect any of the general results of this paper, as both *k*_NM_ and *k*_MN_ are small compared to all other rates, and do not affect the values of the other parameters.

We then compute the ratio *k*_MI_/*k*_NI_ using the best-fit parameters from Table S2. Consistent with the idea that GroEL is much more selective in unfolding only the misfolded states as compared to CYT-19, we find using Eq. (8) *κ* ≈ *k*_NI_/*k*_MI_ ~ (6 – 8)10^−3^, for GroEL in the concentration range between 0.5 - 5*μ*M and 1mM ATP concentration. Compared to the values of (0.0125 – 0.025) that we found for the ribozyme system, this result shows that GroEL is indeed much more effective in discriminating between the misfolded and native states. The steady state yield of the native state as a function of *κ* (Fig. 7), which can be altered by mutations [47], is a highly sensitive function of *κ.* Finally, we also use our model to analyze refolding data on the protein MDH. The excellent fit of data to our model (Fig. 5c, parameters in Table S3) further validates the use of the stochastic 3-state system to describe chaperone assisted refolding of proteins. It is worth noting that sub-stoichiometric quantities of GroEL give substantial yield of the native state, albeit at lower rates [48]. Similar to *T.* ribozyme and Rubisco, the long term native yield of MDH changes as the GroEL concentration changes, indicating that the steady state reached is far from equilibrium. Despite the fundamentally different mechanistic functions, our model provides a unified description of the action of both GroEL and CYT-19.

### Protein and RNA chaperones function far from equilibrium

The quality of our fits in Fig 3, 4, 5 and the consistency of the fitted parameters with previous experimental results, prove the validity of the theory to describe the general behavior of both protein and RNA chaperones. The major conclusion of our work is that the steady state yield of native states *P*(N, ∞) does *not* approach a Boltzmann distribution dictated by equilibrium thermodynamics (see **SI** for more details). The value of *P*(N, ∞) is a function of both the chaperone concentration [C] as well as the ATP concentration [T], as is evident both from Eq. (3) and from the experimental data (Figs. 3, 4, 5). For example, since we calculated the stability of the P5a variant of *T.* ribozyme to be 2.6*k_B_T*, equilibrium thermodynamics would predict a yield 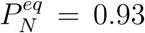. Clearly, this yield is not reached for most concentrations of the chaperone (Fig. 6 b). The steady state yields depend on both ATP and CYT-19, an indication that equilibrium is not reached. Since Δ*G*_NM_ for Rubisco or MDH is not known to the best of our knowledge, we could not do the same analysis on Rubisco or MDH to estimate the native yield predicted by equilibrium thermodynamics. However, the observation that the steady state yield of the native states changes with GroEL concentration (Fig. 5a,c) shows that equilibrium is not attained in the Rubisco–GroEL/MDH systems either. In addition, it is clear from the rate parameters determined in our model that the local detailed balance condition is broken, and that a non-zero probability current is established between any two states in the three state model, further demonstrating that the steady state reached by the chaperone and the system is far from equilibrium (see **SI** for details).

**FIG. 6:**
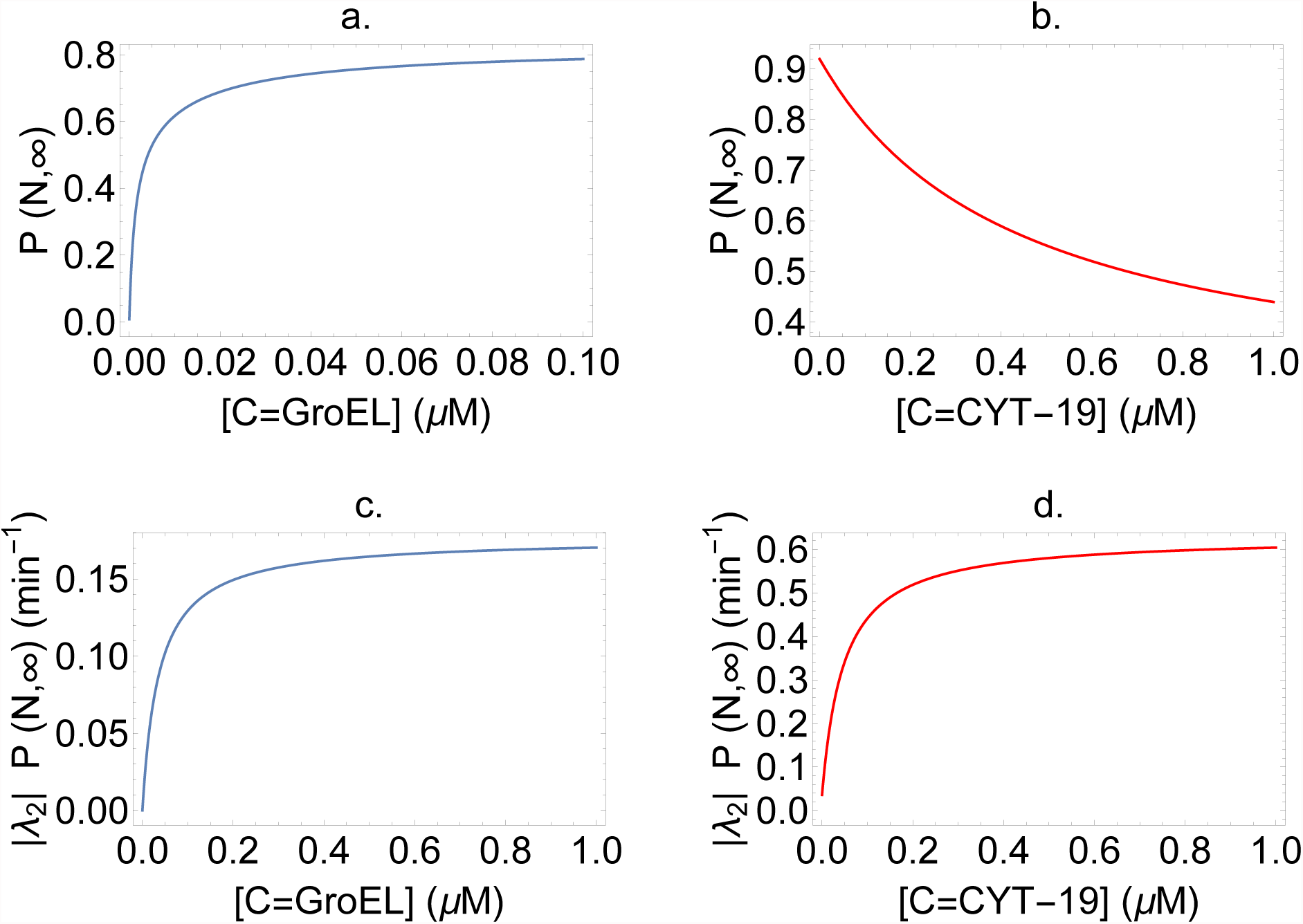
Chaperones maximize the yield of native state per unit time, not the absolute yield. **(a-b)** Steady state yield of the native state of Rubisco (**a**) and ribozyme (**b**), as functions of chaperone concentration. (**c-d**) Native state yield per unit time of Rubisco (**c**) and ribozyme (**d**), as functions of chaperone concentration. The curves in (**a**) and (**c**) were obtained using the best-fit parameters for GroEL–Rubisco system, given in Table S2. The curves in (**b**) and (**d**) have been produced using the best-fit parameters for the mutant P5a ribozyme, given in Table S1. For all the curves, the ATP concentration [T] was set to 1 mM. The qualitative results do not change for other concentrations of ATP.

**FIG. 7:**
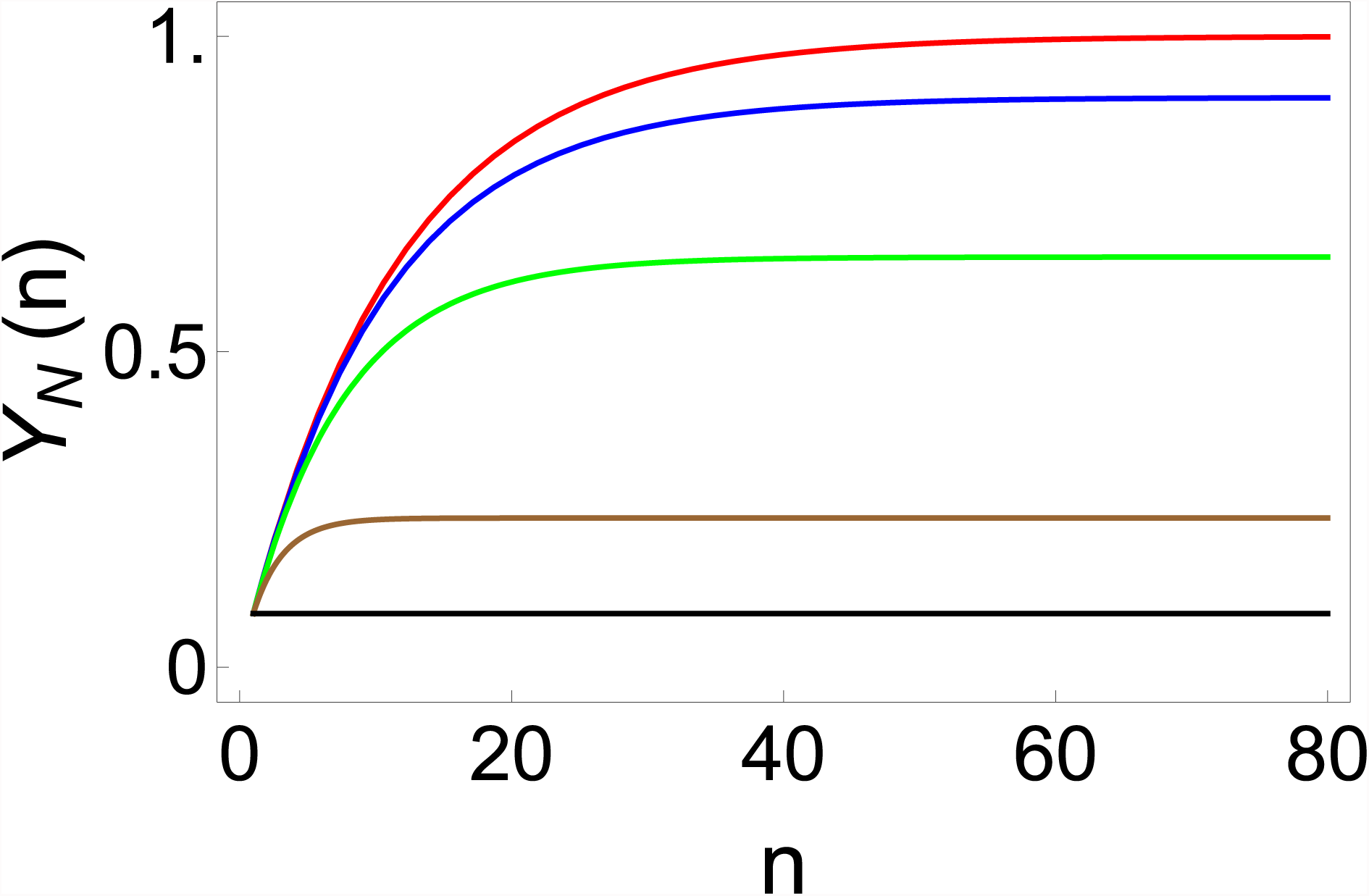
Generalized iterative annealing mechanism for proteins and RNA, showing the effect of varying *κ,* on the yield of the native state. Plot of the yield, *Y_N_*(*n*) (see Eq. 7), as a function of number of cycles *n.* The native fraction in the limit of large *n* therefore depends on *κ,* the efficiency of chaperone recognition of the native state: *κ* = 0 (red), *κ* = 0.01 (blue), *κ* = 0.05 (green), *κ* = 0.3 (brown) and *κ* = 1.0 (black).

### The yield of native state per unit time is maximized by iterative annealing mechanism of the chaperones

Do chaperones maximize the absolute yield of the native states or the rate of their folding? Our theory suggests a general answer to this question, for both RNA and protein chaperones. Using the parameters in Tables S1 and S2, the steady state native yield *P*(N, ∞) is plotted for both Rubisco and *T.* ribozyme in Figs.6 a,b. The figure highlights that increasing the chaperone concentration results in completely opposite behavior of the native yield for Rubisco and *T.* ribozyme. This immediately suggests that the absolute value of the yield is not a quantity that is maximized by chaperones, since increasing chaperone concentration would decrease the native yield as is the case for CYT-19 action on the ribozyme (Fig 6 b). However, as shown in Fig 6 c,d, the rate of increase of the native yield (production of the folded state per unit time) is a monotonically increasing function of the chaperone concentration, reaching saturation values at ~ 1*μM* both for the RNA and the protein. The same plot for MDH is shown in Fig. S1. Given that *in vivo* the number of chaperones is regulated, this remarkable result shows that it is the yield of the native state per unit time that is maximized by the chaperones. This result is further substantiated by the observation that the GroEL concentration *in vivo* is about 5.2 *μM* (there are 1580 14-mer GroEL molecules in a volume of 1 *μm*^3^ in *E. Coli* with the functional unit being the 7-mer). As can be seen from Fig 6 c and Fig. S2, a concentration of 5.2 *μM* is in the saturation region, suggesting that the rate of increase of native yield is maximized.

## Concluding Remarks

With a doubling time of about 2 hours, *Tetrahymena* are some of the fastest multiplying free living eukaryotic cells [49]. Therefore, the viable time scale for *T.* ribozyme folding to the native state, should be on the order of a few hours. Though a large fraction of the ribozyme (1 – Φ with Φ ≈ 0.1) misfolds and stays kinetically trapped over time scales of days *in vitro* [11], experiments show that the addition CYT-19 can accelerate the folding process to a matter of minutes [28]. Surprisingly however, increasing the CYT-19 concentration decreases the final yield of the native states, in stark contrast to GroEL-mediated folding of proteins, where increasing the chaperone concentration increases the native yield at long times.

In this work, we have developed a theoretical model to study the widely contrasting experimental results on protein and RNA chaperones. The representation of assisted folding of protein or RNA using a three–state stochastic model allowed us to explore the concepts of kinetic partitioning factor (Φ), native population at thermo-dynamic equilibrium (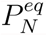), and biologically relevant non-equilibrium steady state yield of the native states (*P*(*N*, ∞)). By using a unified framework to analyze a wide array of data from CYT-19 dependent *T.* ribozyme folding experiments and GroEL mediated Rubisco folding studies, we show unequivocally that thermodynamic equilibrium is not attained in these systems. Rather, the chaperones use energy from ATP hydrolysis to drive the system out of equilibrium, thereby maximizing the rate of yield of native states. This purely non-equilibrium effect is a novel result that is distinct from earlier experimental and theoretical works that have suggested interesting implications for the role of out of equilibrium dynamics on the conformational cycles of chaperones [50] and on the affinity of chaperones for their substrates [51].

The time to increase the native yield from Φ to 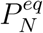 in the absence of chaperones is beyond biologically meaningful time scales. For example, *τ*_*M*→*N*_ ≈ 10^4^ min in the *T.* ribozyme [52]. In other words, this implies that the functions of molecular chaperones is to maximize the yield in a given time, rather than the yield or the folding rate. Assisted folding does not maximize the folding rates either, as erroneously stated in many studies based on probing the efficacy of artificial single mutant (SR1) of GroEL in rescuing substrate proteins. The current work debunks the conclusions of such studies, which have inferred the function of GroEL based on SR1, which is at best a useful model in probing confinement effects on protein folding. Such studies have no relevance to the function of the full chaperonin machinery either *in vitro* or *in vivo.*

Just like error-correcting machineries in cellular processes [53, 54], molecular chaperones utilize energy from ATP hydrolysis to stochastically help fold biomolecules into the functionally active form on biologically meaningful time scales. Our work shows, that at least for this class of stochastic machines, molecular chaperones have evolved to maximize the rate of native-state production per unit time, even if accompanied by lavish consumption of ATP. Our results provide a unified framework for the role of chaperones in both protein and RNA folding, and may have important implications in our understanding of both protein and RNA homeostasis [55, 56].

## Acknowledgements

Much of this work was done while SC and DT were at the Korea Institute of Advanced Study in 2013. We are grateful to the National Science Foundation through grant CHE 16-36424 and the Collie-Welch Chair for supporting this work.

## I. SOLUTION OF THE 3-STATE MODEL

Denoting by 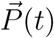 a column vector with components *P*(*i, t*), Eq. 2 of the main text can be cast into a matrix form: 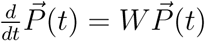 where the rate matrix *W* is

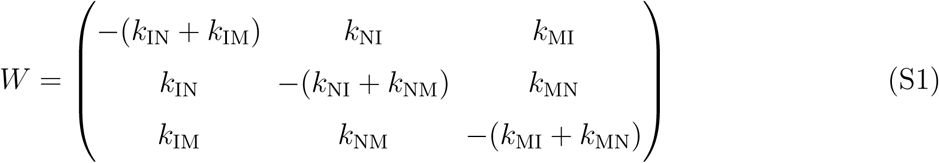

For the rate matrix in Eq. (S1), 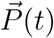 is given by,

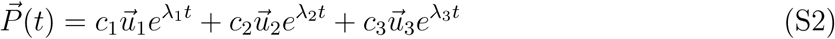

where λ_1,_ λ_2_ and λ_3_ are the eigenvalues of W and 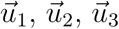 are the corresponding eigenvectors. The first eigenvalue λ_1_ is equal to 0 while λ_2_, λ_3_ < 0 [1]. Since λ_1_ = 0 (and the other two eigenvalues are negative), 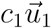 represents the steady–state (*t* → ∞) solution of Eq. S2. If |λ_2_| ≪ |λ_3_|, Eq. S2 effectively describes a single exponential relaxation to the steady-state, with |λ_2_| the observed rate for the relaxation, i.e., |λ_2_| ≈ *k_obs_.* The coefficients *c*_1_*, c*_2_ and *c*_3_ are constants determined from the initial conditions—the fraction of substrate in I, M and N at time *t* = 0. In all the experiments analyzed later, three types of initial conditions arise: all the substrate begins in state M (*P*(M,0)=1), all the substrate begins in state N (*P*(N,0)=1) and the substrate is in a mixture of states. The last initial condition was needed only for analyzing some of the ribozyme data (Figs. 3 and 4), and for these cases, *P*(N,0) was obtained directly from the experimental data at *t* = 0 and P(M,0) was set to 1 – *P*(N, 0).

## II. THE LONG TIME STEADY STATE IS FAR FROM EQUILIBRIUM

To assess whether the long term steady state of Eq. S2 is an equilibrium or non-equilibrium solution, we calculate the probability current between any two states of the model. This current *J* is the same between any two states of our 3-state model, and is defined as:

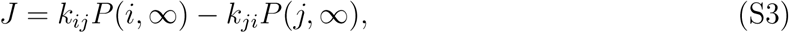

where *i, j* =I,M,N. Using Eq. S2 with *t* → ∞, the current *J* is given by,

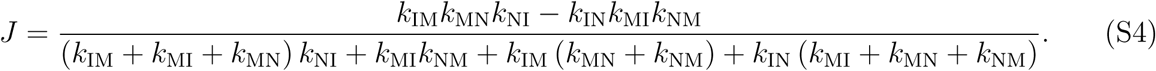

The steady state reached is out of equilibrium, as in the long time limit a non-zero (but constant) probability current *J* exists between any two states of the system. As is evident from Eq. (S4), only when both *k*_NI_ = 0 and *k*_MI_ = 0, which is realized either when [C]=0 or [T]=0 or both, the current becomes zero which is characteristic of equilibrium.

## III. PARAMETER ESTIMATES FOR RNA AND PROTEINS

To analyze the fraction of native substrate as a function of time, we fit Eq. S2 to experimental data using a custom written non-linear least squares method in Mathematica [2]. The least squares method of minimizing *χ*^2^ values is equivalent to maximizing a log-likelihood function, with the assumption that errors in the mean fraction of native substrate are Gaussian-distributed. As can be seen from Tables S1–S3, a number of parameters could not be uniquely identified from the fits. To find ranges of these parameters over which the fits did not appreciably change, we varied each parameter individually around the best fit value keeping all other parameters fixed, and computed the change in the minimum *χ*^2^. We reported parameter ranges that allowed the *χ*^2^ value to increase by 1 from the minimum. Quantitatively, this means that a parameter resulting in an increase of 1 for the *χ*^2^ is exp(–1/2) ~ 60% as likely to be correct as the best fit parameter value.

This is physically reasonable since the chaperone is expected to bind more efficiently to misfolded protein rather than native protein, and was found to be the case for the unconstrained fit for Rubisco shown in Table S2.

## IV. YIELD PER UNIT TIME OF NATIVE MDH

Fig. S1 shows the yield per unit time of the protein MDH as a function of GroEL concentration. Similar to Rubisco and T.ribozyme analyzed in the main text, this is an increasing function of chaperone concentration. The curve appears to saturate at GroEL concentration exceeding approximately 2*μM.* This finding and the observation that the *in vivo* GroEL concentration is approximately 2.6 *μM* further supports our prediction GroEL maximizes the native yield per unit time and not the folding rate.

## V. PREDICTIONS FOR POSSIBLE MUTATIONS

Having obtained the best fit parameters for the ribozymes and proteins (Table S1, S2 and S3), we can now modify the rates of some of the important parameters, to predict the outcome of mutations that could conceivably be performed. The results, shown in Fig. S2 for the WT ribozyme and Fig. S3 for MDH, suggest that the most sensitive mutation would be one that changes the binding of the chaperone to native ribozyme or protein. Interestingly, our analysis predicts that other possible mutations (that would change ATP hydrolysis rates or binding rates of chaperone to the misfolded ribozyme/protein) would hardly change the final yield of native states.

---

**TABLE S1:**
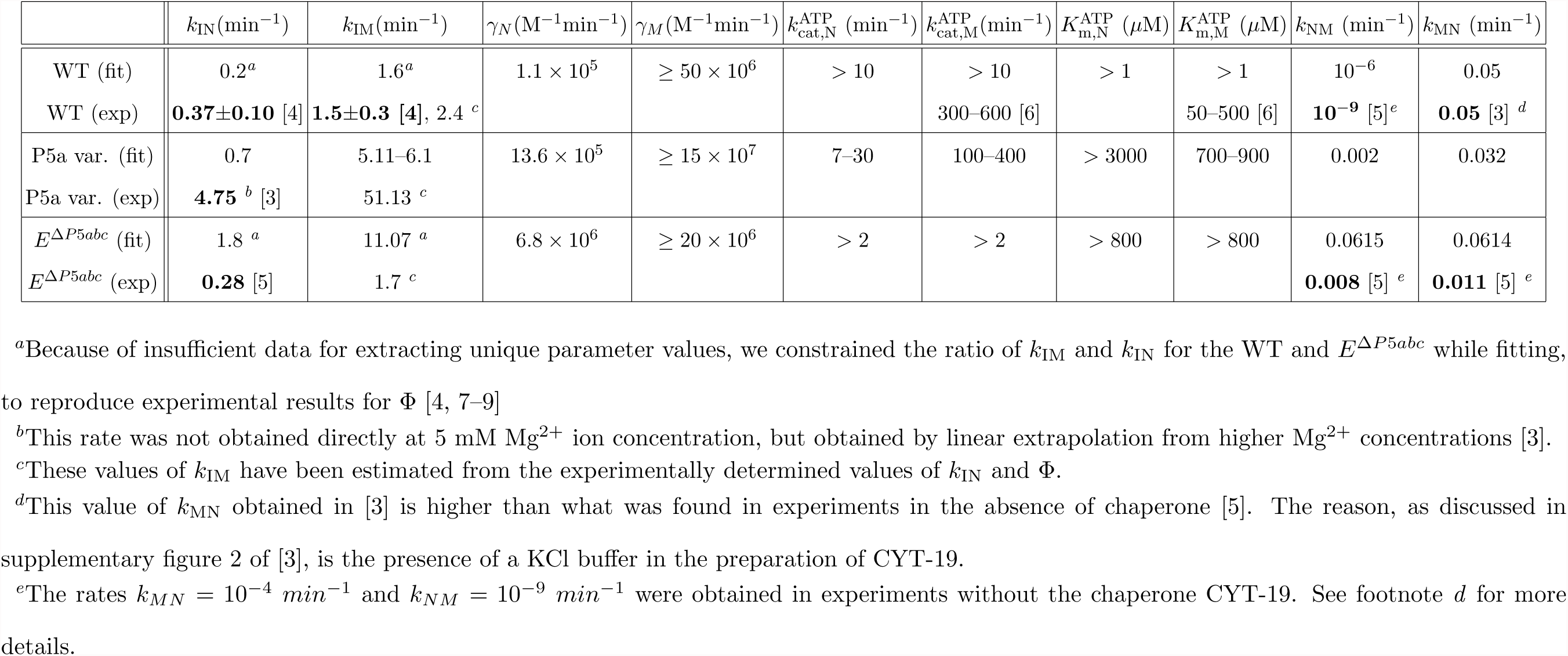
Best-fit parameters extracted by fitting Eq. S2 to the experimental data [3] obtained at 25 °C and 1 mM (WT), 5 mM (P5a and *E*^ΔP5*abc*^) Mg^2+^ ion concentrations. For comparison, we list the corresponding rates from direct experimental measurements (in bold) and the indirect (details in the footnotes below) estimates. The experimental rates were determined at 25 °C, but with different Mg^2+^ ion concentrations, 10 mM in [4] and 10–50 mM in [5].

**TABLE S2:** Best-fit parameters determined by fitting Eq. S2 to experimental data on GroEL-mediated folding of Rubisco [10].

| | $k_{\text{IN}} (\text{min}^{-1})$ | $k_{\text{IM}} (\text{min}^{-1})$ | $\gamma_N (\text{M}^{-1} \text{min}^{-1})$ | $\gamma_M (\text{M}^{-1} \text{min}^{-1})$ | $k_{\text{cat},N}^{\text{ATP}} (\text{min}^{-1})$ | $k_{\text{cat},M}^{\text{ATP}} (\text{min}^{-1})$ | $K_{\text{m},N}^{\text{ATP}} (\mu\text{M})$ | $K_{\text{m},M}^{\text{ATP}} (\mu\text{M})$ | $k_{\text{NM}} (\text{min}^{-1})$ | $k_{\text{MN}} (\text{min}^{-1})$ |
| --- | --- | --- | --- | --- | --- | --- | --- | --- | --- | --- |
| fit | 0.25 | 20.36 | $27 \times 10^5$ | $389 \times 10^6$ | 0.02 | 48 – 52 | < 40 | 60 – 145 | $10^{-6} - 2 \times 10^{-4}$ | 0.0094 - 0.01 |

**TABLE S3:** Best-fit parameters determined by fitting Eq. S2 to experimental data on GroEL-mediated folding of MDH.

| | $k_{IN} (\text{min}^{-1})$ | $k_{IM} (\text{min}^{-1})$ | $\gamma_N (\text{M}^{-1} \text{min}^{-1})$ | $\gamma_M (\text{M}^{-1} \text{min}^{-1})$ | $k_{cat,N}^{\text{ATP}} (\text{min}^{-1})$ | $k_{cat,M}^{\text{ATP}} (\text{min}^{-1})$ | $K_{m,N}^{\text{ATP}} (\mu\text{M})$ | $K_{m,M}^{\text{ATP}} (\mu\text{M})$ | $k_{NM} (\text{min}^{-1})$ | $k_{MN} (\text{min}^{-1})$ |
| --- | --- | --- | --- | --- | --- | --- | --- | --- | --- | --- |
| fit | 0.366 | 0.371 | $< 10^{3c}$ | $1.7 \times 10^{6c}$ | $> 0.1^a$ | $> 20^a$ | $< 2000^b$ | $< 2000^b$ | $7.78 \times 10^{-3}$ | 0.025 |

<sup>a</sup>Because of insufficient data for extracting unique parameter values, the $k_{cat}$ values were constrained to be less than $5000 \text{ min}^{-1}$ , since most enzymes fall in this range [11]
<sup>b</sup>Because of insufficient data for extracting unique parameter values, the $K_m$ values were constrained to be less than $10000 \mu\text{M}$ , since most enzymes fall in this range [11]
<sup>c</sup>Because of insufficient data for extracting unique parameter values, the additional constraint of $\gamma_M > \gamma_N$ was maintained while fitting to data. This is physically reasonable since the chaperone is expected to bind more efficiently to misfolded protein rather than native protein, and was found to be the case for the unconstrained fit for Rubisco shown in Table S2.

**FIG. S1:**
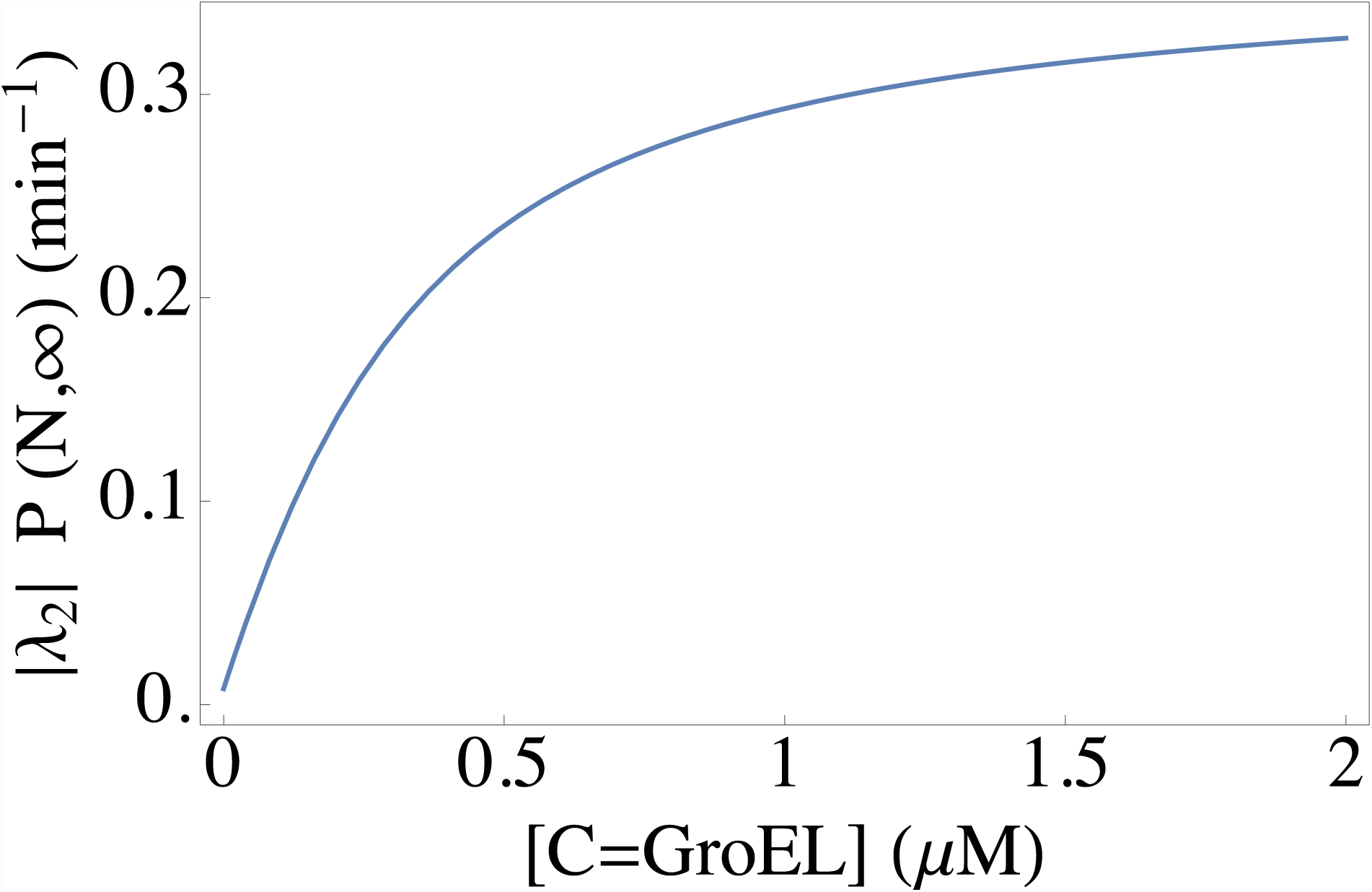
The yield of native MDH per unit time as a function of GroEL concentration (7 or 14-mers of GroEL). Concentration of MDH was 0.5*μM.*

**FIG. S2:**
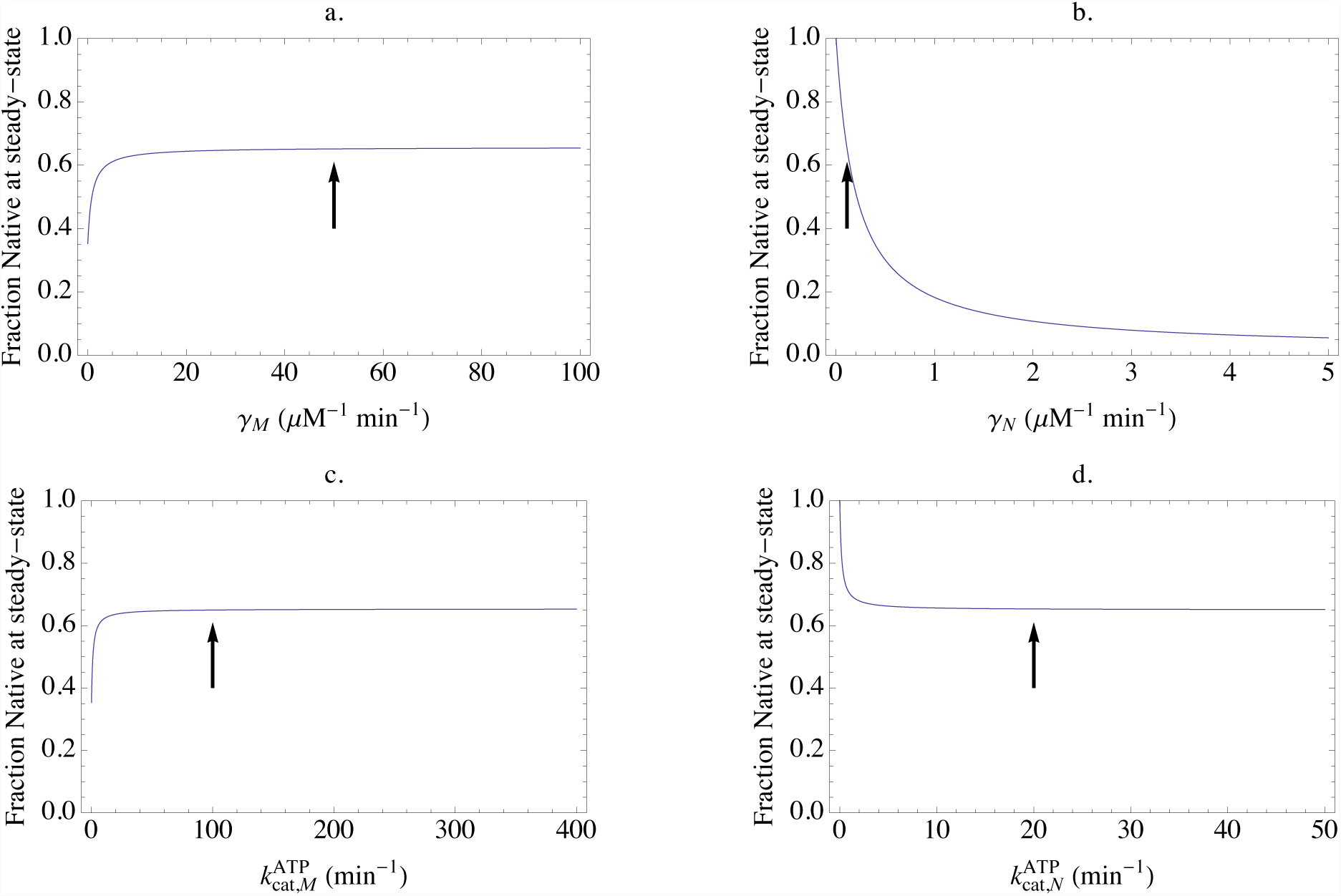
Effect of possible mutations on the final yield of native states in the WT ribozyme–CYT-19 complex. Parameters were fixed to the best-fit results with [C]=1*μ*M, [T]=2000*μ*M and only γ*_M_* (panel a.), γ*_N_* (panel b.), 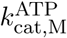 (panel c.) and 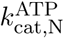 (panel d.) were varied to observe the effect on the fraction of native states. Arrows indicate the position of the best fit value around which the parameter is being varied.

**FIG. S3:**
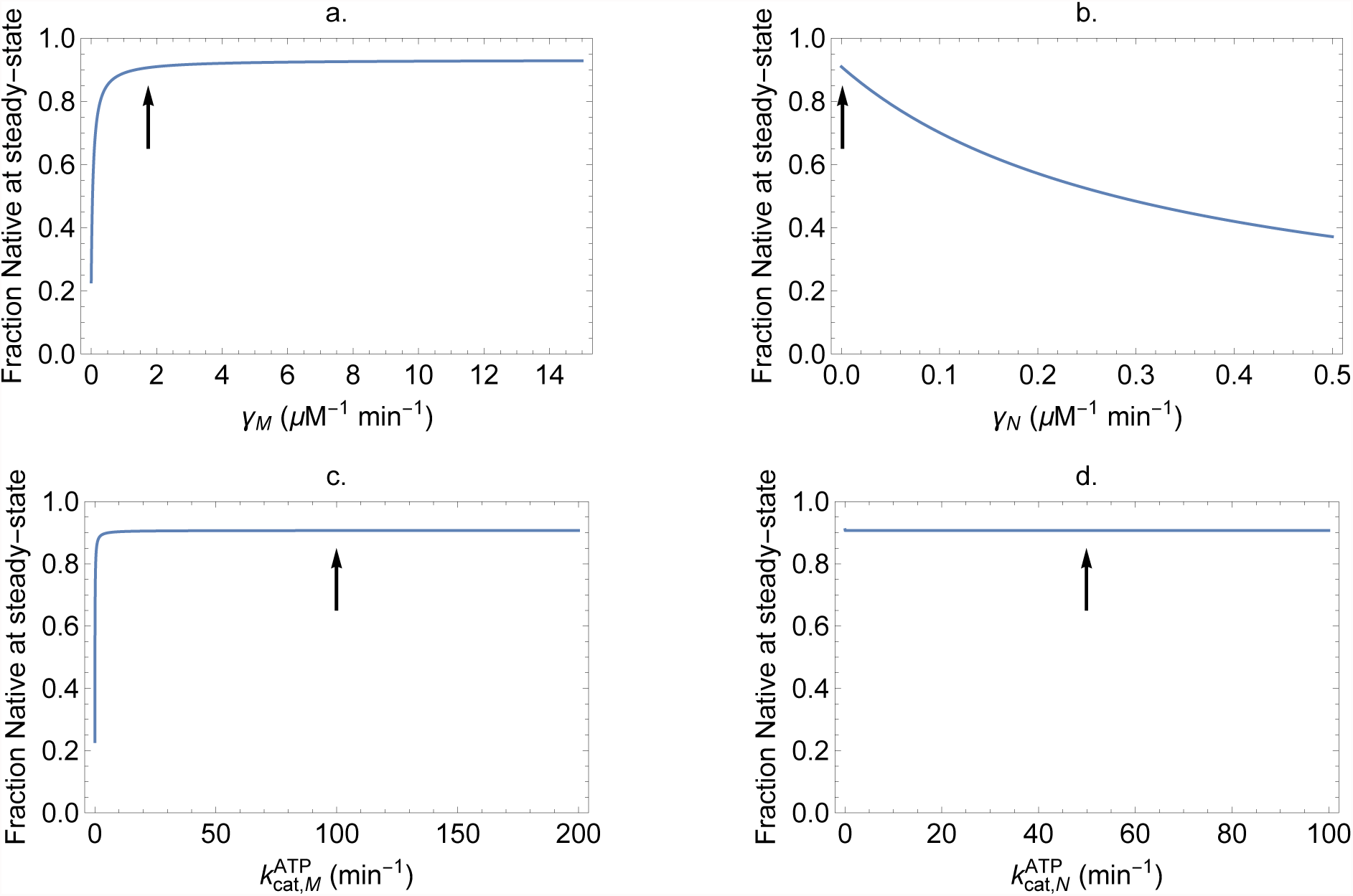
Effect of possible mutations on the final yield of native states in the MDH–GroEL complex. Parameters were fixed to the best-fit results with [C]=1*μ*M, [T]=2000*μ*M and only γ*_M_* (panel a.), γ*_N_* (panel b.), 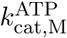 (panel c.) and 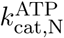 (panel d.) were varied to observe the effect on the fraction of native states. Arrows indicate the position of the best fit value around which the parameter is being varied.

